# Metabolic flux fingerprinting differentiates planktonic and biofilm states of *Pseudomonas aeruginosa* and *Staphylococcus aureus*

**DOI:** 10.1101/2020.07.15.203828

**Authors:** Mads Lichtenberg, Kasper Nørskov Kragh, Blaine Fritz, Julius Bier-Kirkegaard, Thomas Bjarnsholt

## Abstract

The challenges of defining the biofilm phenotype has been clear for decades. Many biomarkers for biofilm are known, but methods for identifying these are often invasive and/or complicated. These methods often rely on disrupting the biofilm matrix or examining virulence factors and compounds, which may only be expressed under certain conditions.

We used microcalorimetric measurements of metabolic energy release to investigate whether unchallenged, planktonic *Pseudomonas aeruginosa* displayed differences in metabolism compared to surface-bound and non-attached biofilms.

The pattern of energy release observed in the recorded microcalorimetric thermograms clearly depended on growth state, though the total energy expenditure was not different between growth states. To characterize these differences, we developed a classification pipeline utilizing machine learning algorithms to classify growth state, based on the observed patterns of energy release. With this approach, we could with high accuracy detect the growth form of blinded samples. To challenge the algorithm, we attempted to limit the amount of training data. By training the algorithm with only a single data point from each growth form, we obtained a mean accuracy of 90.5% using two principal components. Further validation of the classification pipeline showed that the approach was not limited to *P. aeruginosa* but could also be used for detection of gram-positive *Staphylococcus aureus* biofilm. We propose that microcalorimetric measurements, in combination with this new quantitative framework, can be used as a non-invasive biomarker to detect the presence of biofilm.

These results could have a significant potential in clinical settings where the detection of biofilms in infections often means a different outcome and treatment regime for the patient.

## Introduction

Bacteria residing in biofilms are thought to be phenotypically distinct from their planktonic counterparts. Many publications have identified specific biomarkers to characterize this distinction, such as differential gene expression (Folsom *et al*., 2010), secretion of extracellular polymers (Costerton *et al*., 1995; Goltermann and Tolker-Nielsen, 2017), virulence factors (Hauser, 2011), metabolism (Solokhina *et al*., 2017) and increased tolerance towards antibiotics and host response (Bjarnsholt *et al*., 2013; Ciofu and Tolker-Nielsen, 2019). However, physical aggregation (observed by microscopy) and increased antibiotic tolerance are the most dominant and consistent of these biofilm-associated phenotypes. Traditionally, biofilm research has focused on surface-attached biofilms, but there is an increasing focus on embedded and non-attached biofilm aggregates (Secor *et al*., 2018; Alhede *et al*., 2011; Kragh *et al*., 2016). In the majority of biofilm-related infections, bacteria are found as non-attached aggregates embedded in host material, such as slough or mucus (Bjarnsholt *et al*., 2013). The difference in growth geometry between planktonic bacteria, surface-attached, and non-attached biofilms may result in distinctive microenvironments (Stewart and Franklin, 2008; Stewart *et al*., 2016; 2019), which could differentially impact the metabolic activity and number of bacteria aggregating (Sønderholm *et al*., 2018) and vice versa.

The increased tolerance of biofilms toward antibiotics is frequently attributed to a lower metabolic rate (Kolpen *et al*., 2017), upregulated efflux pumps (Frimodt-Møller *et al*., 2018; Bartell *et al*., 2019), protection by matrix components (Tseng *et al*., 2013; Cao *et al*., 2016) or SOS responses (Nguyen *et al*., 2011; Bernier *et al*., 2013), and is determined by direct exposure to antibiotics and measure of killing either by plating and enumeration of surviving colony forming units (CFU) or live/dead staining. The CFU method, however, can be slow and prone to biases, such as counting aggregates as single CFUs or false negatives due to the induction of viable but non-culturable state of the bacteria (Kvich *et al*., 2019). Similarly, live/dead staining may overestimate the proportion of dead cells, due to the binding of propidium iodide to eDNA (Rosenberg *et al*., 2019).

A measurable biomarker is needed in order to move the field forward and improve our understanding of the difference between bacteria living as single-cells or in biofilms (Jefferson, 2004).

Non-invasive measurements of activity in biofilms are not trivial, as many current methods rely on disrupting the biofilm and altering the chemical microenvironment, though some genetic (Poulsen *et al*., 1993; Kragh *et al*., 2014) and chemical reporters do exist (Corte *et al*., 2019). Most of these methods, however, only cover part of the metabolic output linked to, e.g., growth or production of molecules produced under certain conditions (Whooley and McLoughlin, 1982; Poulsen *et al*., 1993). Since metabolic activity seems to play a distinguishing role between bacteria in planktonic-and biofilm states, a fine-scale measurement of energy release could be a biomarker of interest.

Microcalorimetry has been used in material research for decades, but is gaining interest in life sciences (Braissant *et al*., 2015; Baldoni *et al*., 2009; Butini *et al*., 2019; Tellapragda *et al*., 2020). This technique records energy release in the form of heat flow, providing a real-time measurement of the population-level metabolic activity in a sample (Braissant *et al*., 2015; Wadsö *et al*., 2017).

In the present study, we examined the heat flow of planktonic bacteria (PC) and biofilm-associated bacteria (Fig. 1) using non-invasive, isothermal microcalorimetry. Due to the increased focus on non-attached biofilm aggregates, we also compared the metabolism of non-attached aggregates embedded in alginate beads (AB) with surface-attached biofilms (SB) (Alhede *et al*., 2011; Sønderholm *et al*., 2017b).

**Figure 1.**
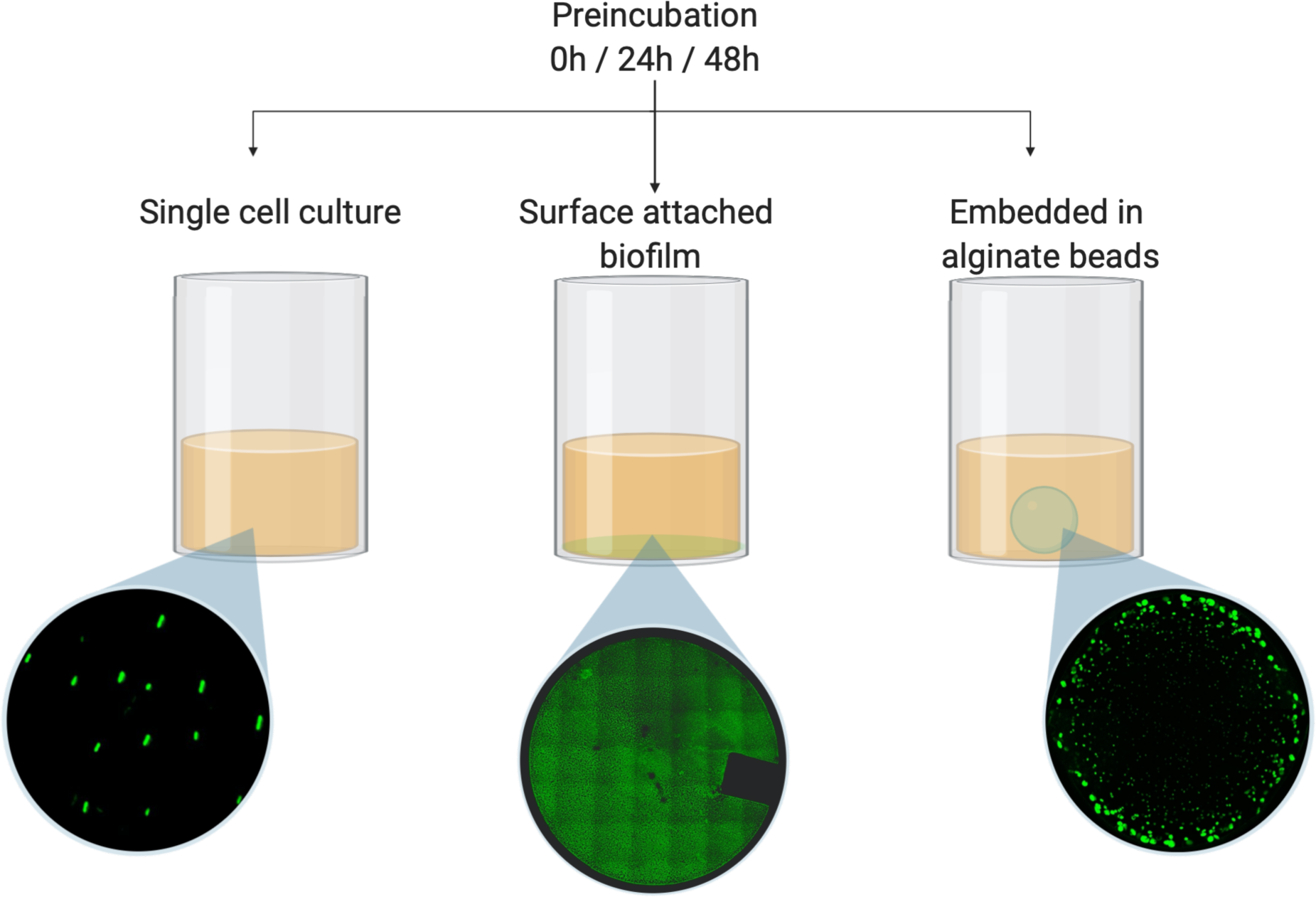
Schematic drawing of the experimental conditions. *Pseudomonas aeruginosa* or *Staphylococcus aureus* was either grown as planktonic culture (PC), as surface-attached biofilm (SB), or as aggregates embedded in alginate beads (AB). The three models were measured in an isothermal microcalorimeter with 0h, 24h or 48h preincubation (surface-attached biofilm = only 24h and 48h preincubation). Microscope images were acquired on a confocal laser scanning microscope (Zeiss 880LSM) using a strain of PAO1 with constitutively expressed GFP.

We aimed to investigate whether bacteria growing in biofilms displayed distinct metabolic features compared to the planktonic state.

## Results

### Metabolic activity of alginate bead biofilms, surface-attached biofilms and planktonic culture

After equilibration of the thermal signal (∼30 min), all three models containing *P. aeruginosa* showed signal increases. Thermograms from all models showed 2-3 peaks in heat flow, but the position and magnitude of the peaks varied between models.

The accumulated energy in each well was 2.19 J ± 0.17 (mean ± SD) and did not vary within or between conditions (p>0.05; One-way ANOVA test). In two follow-up experiments, we tested whether the cease in heat flow was related to electron acceptor or -donor limitation by 1) opening the wells to let in new atmospheric air after the thermogram had reached zero after which we observed a second peak, and 2) by adding 10mM NO_3_^-^. When this was done, the total energy increased by ∼35% for the alginate embedded biofilm, and we concluded that, under the specifications used for these experiments, the heat flow reached zero due to electron acceptor limitation rather than carbon source limitation.

The recorded thermograms displayed some variation in temporal dynamics as well as in the magnitude of the heat flow within experiments. We tested possible explanations and inoculated increasing amounts of bacteria into alginate beads and as planktonic cells to test if the variation could be explained by bacterial concentration. These variations resulted in both lateral and horizontal shifts in the position of peaks (Fig. S3) as also shown for other microbes previously (Braissant *et al*., 2015; Wadsö *et al*., 2017).

The number of viable bacteria in each well were generally not statistically different between models or preincubation times neither at the start or at the end of experiments (p>0.05; Tukey’s multiple comparison test), though the 24h and 48h preincubated alginate beads contained more bacteria than the other models at the start of the experiment but not at the end (p<0.01; Tukeys multiple comparison test) (Fig. S1).

**Figure 2.**
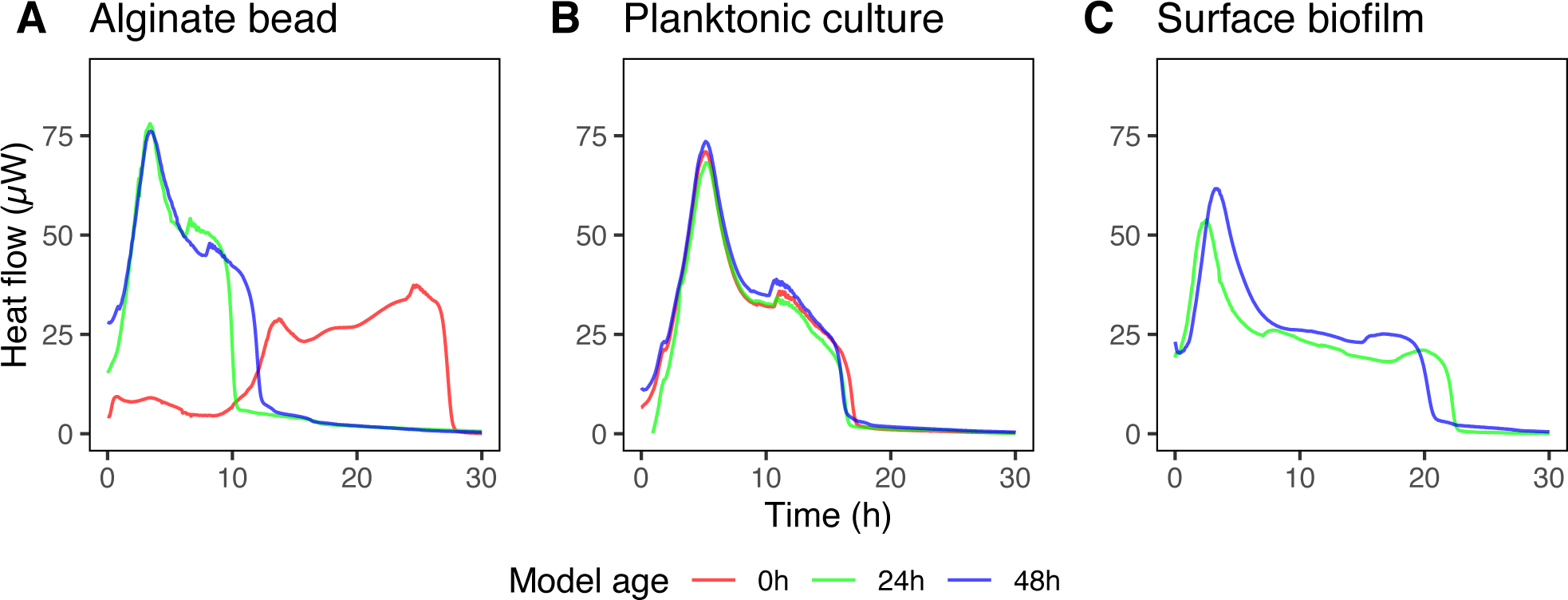
Example of thermograms for *Pseudomonas aeruginosa* grown A) in alginate beads, B) as planktonic culture, and C) as a surface biofilm. Colors correspond to preincubation times of 0h (red), 24h (green) and 48h (blue).

### Numerical model of metabolic output

To investigate the systematic differences in the thermograms between the different growth-models, we fitted the experimental data to a mathematical model, assuming that energy release was non-linearly correlated to bacterial concentration (competition for space) and O_2_ availability. Metabolic curves of *P. aeruginosa* were then calculated based on the theoretical bacterial concentration curves. To simplify such a model, we ignored the differences in spatial organization and modeled the system as a well-mixed suspension, which is only valid for the planktonic culture at t=0. The parameters for the biofilm growth forms are thus to be seen as *effective*. For mathematical details on the model used, see Supporting Information.

We solved the coupled system of differential equations numerically to theoretical curves of metabolism (Fig. 3). We expect that many of the parameters are shared between the datasets, since we used the same species and strain of bacteria. We divided the data into four categories: AB inoculated at t = 0 hours, AB at t = 24 and 48 hours, PC, and SB and performed simultaneous fits on the four different growth-forms, forcing specific parameters to be shared (Table 1). The model represented the data well, although it did not describe the smaller secondary peaks observed. These may be a product of switching to electron acceptors with lower energy yields such as NO_3_^-^ or pyruvate which were not included in this model (see Discussion). The effective parameters obtained from the fits are shown in Table 1.

**Figure 3.**
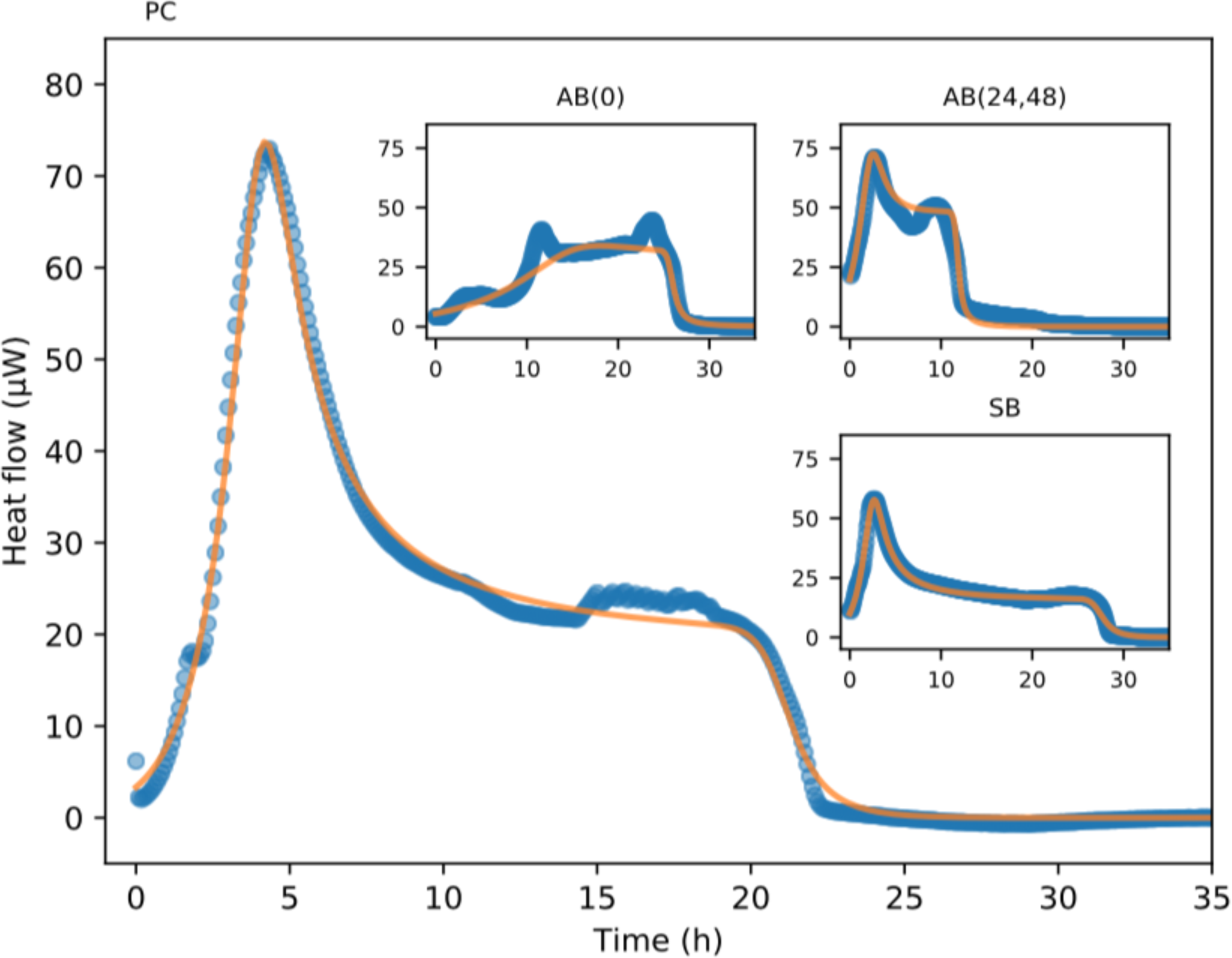
Numerical model of metabolic output of *Pseudomonas aeruginosa* with non-linear dependencies on the concentration of bacteria and O_2_. Some parameters were kept constant between models, while others were allowed to be fitted (see Table 1 and main text). The main graph shows planktonic culture data (PC). Insets show fit to alginate bead biofilm (AB) and surface biofilm (SB). Blue circles represent empirical data, while orange lines are the fitted numerical model. R2 values are shown in Table 1.

**Table 1:** Fixed parameters among all fits in the numerical model

| Yield<br>$1/Y \text{ (h}^{-1}\text{)}$ | Unit conversion<br>$a \text{ (}\mu\text{W)}$ | Non-growth $\text{O}_2$ consumption<br>$\varepsilon$ | Exponent<br>$\gamma_\rho$ | $\text{O}_2$ non-linearity<br>$k_c$ | Asymmetry exponent<br>$\gamma_c$ |
| --- | --- | --- | --- | --- | --- |
| 0.13 | 6.2 | 0.0020 | 3.5 | 36 | 2.7 |

*Variable parameters between fits in the numerical model*
| | Growth rate<br>$\alpha \text{ (h}^{-1}\text{)}$ | Death rate<br>$\beta \text{ (h}^{-1}\text{)}$ | Initial concentration<br>$b(0)$ | Non-linearity<br>$K_a$ | Goodness of fit<br>$R^2$ |
| --- | --- | --- | --- | --- | --- |
| AB (0) | 0.28 | 0.14 | 0.064 | 1.40 | 0.920 |
| AB (24, 48) | 0.96 | 0.25 | 0.23 | 0.65 | 0.970 |
| PC (0, 24, 48) | 0.89 | 0.045 | 0.039 | 0.64 | 0.993 |
| SB (24, 48) | 0.87 | 0.049 | 0.12 | 0.82 | 0.990 |

Evidently, the growth-model influences physiological parameters where, e.g. bacteria pre-inoculated in alginate beads showed higher death rates compared to when they were grown as planktonic culture and as a surface biofilm. The growth rate of AB(0) was lower than all the remaining conditions and also displayed the most substantial growth non-linearity evident from the different shape of the thermogram compared to the other growth-forms that were pre-incubated.

### Cluster analysis

To assess whether thermograms of *P. aeruginosa* grown in different models showed distinct clustering, we made principal component analysis (PCA) with no scaling, which showed clear separation of the different models across the two first principal components (Fig. S2). Thermograms were recorded in models with varying times of preincubation of 0, 24, and 48 hours. In most cases, the model seemed to separate the thermograms as opposed to the preincubation time. However, the newly inoculated alginate beads (AB0) clustered independently of all other models and preincubation times primarily along the first principal component, which accounted for 55.4 % of the total variance. Interestingly, it seems that the switch from single-cell lifestyle to biofilm is explained by the second principal component, which accounted for 22.9 % of the total variance.

### Machine learning algorithm - Framework for detection of growth mode

To demonstrate the feasibility of using the obtained signals to distinguish the mode of growth, we applied machine learning algorithms to classify samples.

Again, we divided the data into four categories: AB inoculated at t = 0 hours, AB at t = 24 and 48 hours, PC, and SB. Using a standard 50 % train-test split on the data, we employed a Gaussian Process with radial basis kernels and, with limited modifications, immediately obtained 100 % classification accuracy on the data. This demonstrated the quality of the data for segmenting the mode of growth. To challenge this, we tried to limit the training data and see how well a Gaussian Process could do by only training with a single data point of each category. A major hindrance of using just a single data point of each category is that there is no information of data variation. To compensate for this, we augmented the data point with warped versions of itself. In particular, we employed smooth spatio-temporal warping kernels that distort the original signals in amounts that resemble the variations found in the original datasets (for further detail see SI).

Using just a single data point from each growth-form, we achieved a mean accuracy of 90.5 % using two principal components (classification accuracy for each category was 0.95 (AB0), 0.87 (AB24,48), 0.88 (PC), and 0.94 (SB)). When we expanded to 9 data points from each growth form, the accuracy increased to 98 %, on average. Using 20 % of the entire dataset, we obtained 100 % accuracy on the data set directly.

**Figure 4.**
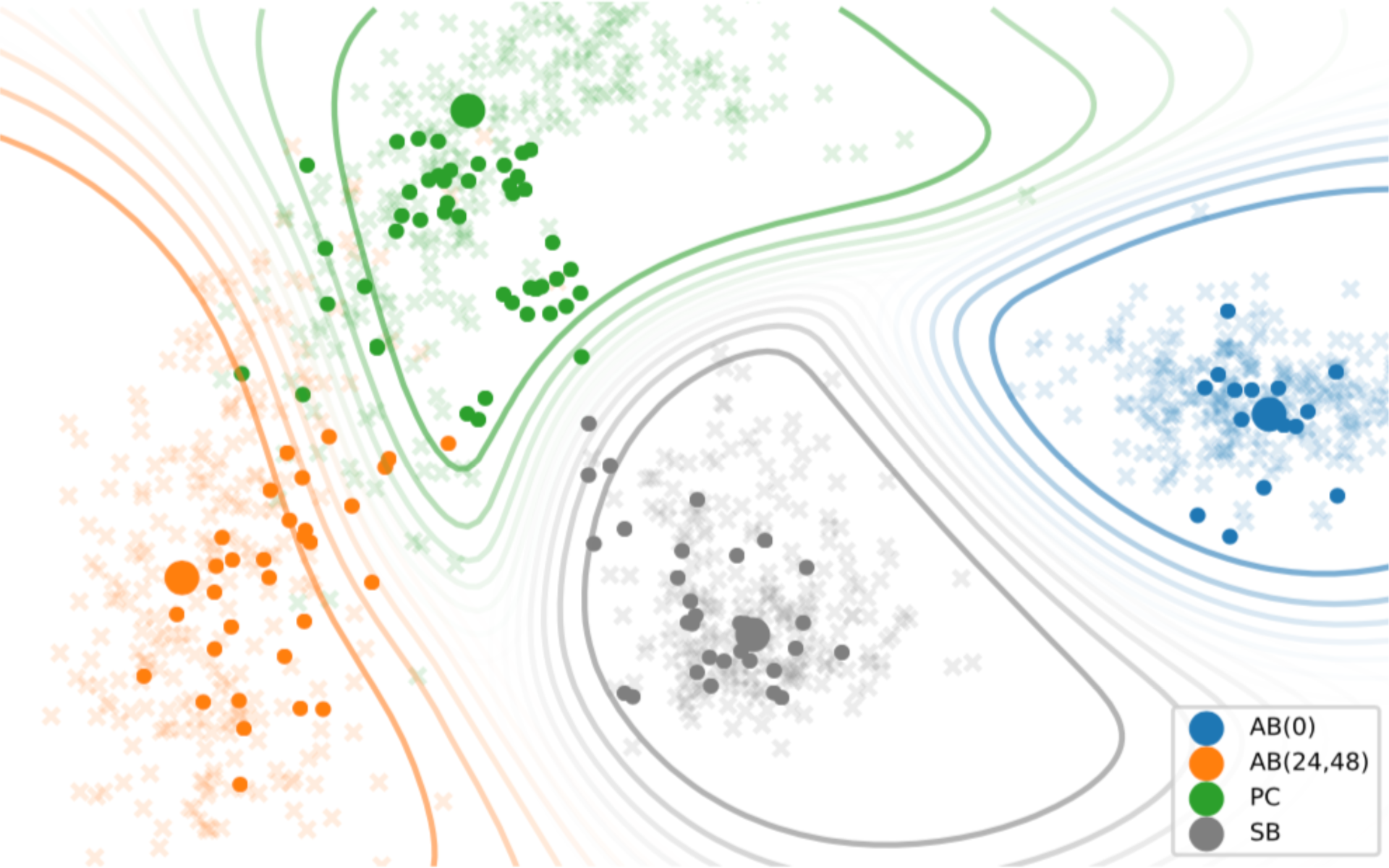
Machine learning classification algorithm developed for thermograms of *Pseudomonas aeruginosa* grown in alginate beads (AB), in planktonic culture (PC), and as surface biofilms (SB) with various preincubation times (0, 24, 48 hours). Large, solid circles represent the data points that were used to train the algorithm; small, solid circles represent the empirical data, and crosses represent the augmented data created by the warping functions. Contours show different categories where the gradient colors of the contours correspond to probability.

### Validation of classification approach

To further validate our method for classifying planktonic from biofilm growing bacteria, we tested the method on the gram-positive coccoid bacterium *Staphylococcus aureus*. Metabolic thermograms showed overall different shapes than displayed by *P. aeruginosa* (Fig. S6). We again tried to classify the samples using the machine learning algorithm using the same growth-form classifications as for *P. aeruginosa* (Fig. 5). Here, the clusters were slightly different from what was observed for *P. aeruginosa*. AB24, and −48 grouped independent of all other conditions but closer to AB0 than seen for *P. aeruginosa*. Surprisingly, the planktonic cultures grouped together with the 24 h preincubated surface biofilm while the 48 hour preincubated surface biofilm grouped closer with the other “biofilm-states”. The average classification accuracy for each category was 0.95 (AB0), 0.96 (AB24,48), 0.56 (PC), and 0.39 (SB).

**Figure 5.**
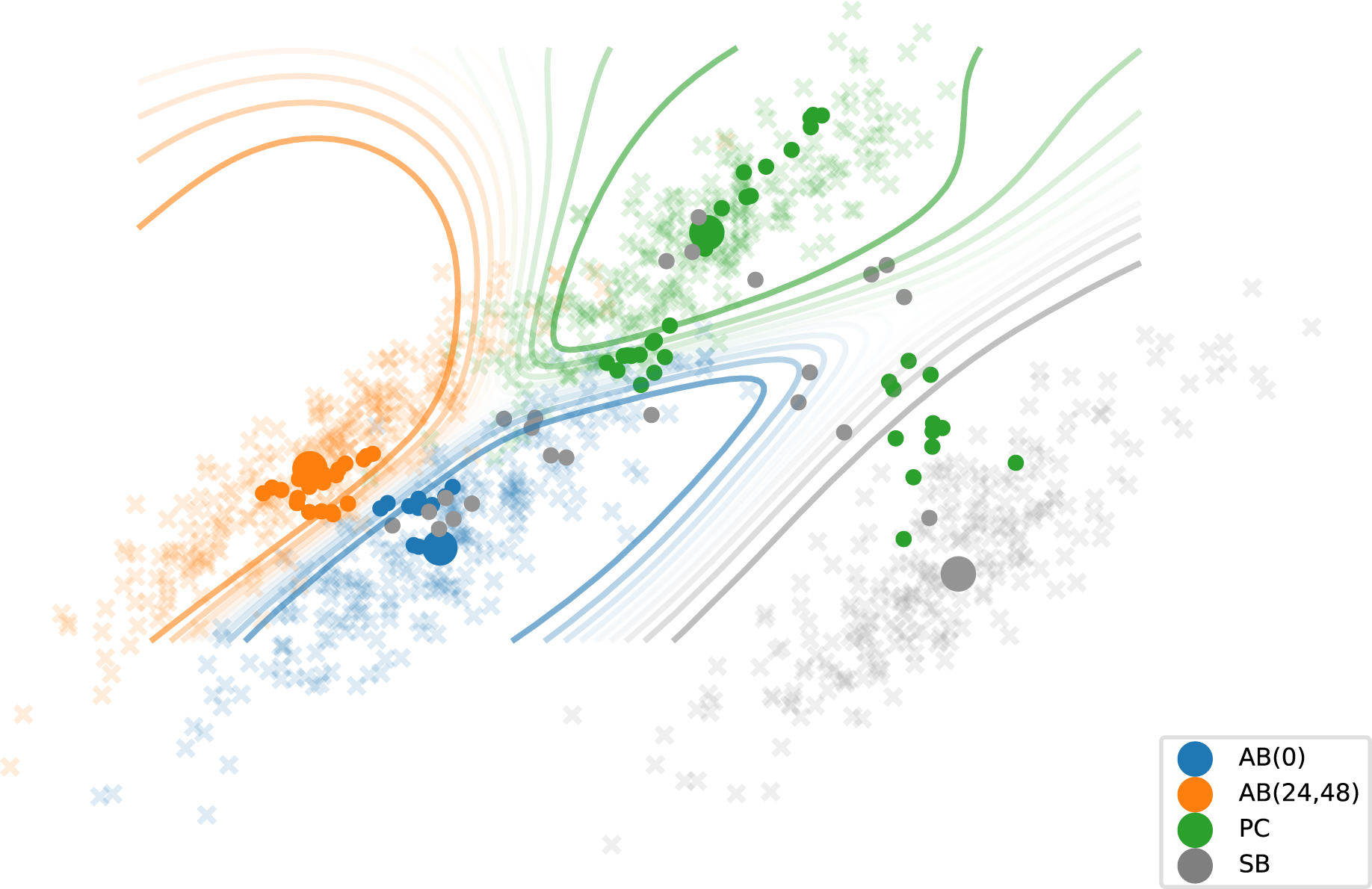
Machine learning classification algorithm applied on thermograms of *Staphylococcus aureus* grown in alginate beads (AB), in planktonic culture (PC), and as surface biofilms (SB) with various preincubation times (0, 24, 48 hours). Large, solid circles represent the data points that were used to train the algorithm; small, solid circles represent the empirical data, and crosses represent the augmented data created by the warping functions. Contours show different categories where the gradient colors of the contours correspond to probability.

## Discussion

Since the first descriptions of biofilm in modern microbiology around 40 years ago (Geesey *et al*., 1977; Høiby, 1977; Jendresen and Glantz, 1981; McCoy *et al*., 1981), considerable scientific research has focused on discovering which factors define a biofilm. Many biomarkers have been proposed, such as the irreversible attachment to a surface, deployment of quorum sensing (QS) systems, overall altered transcription profile, and increased antibiotic tolerance. However, the only consistent “biofilm-phenotypes” are physical aggregation (attached as well as non-attached) and increased antibiotic tolerance. The mechanism that governs this increased tolerance is continuously debated and has, for some types of antibiotics, been linked to protection by matrix components (Tseng *et al*., 2013; Cao *et al*., 2016; Goltermann and Tolker-Nielsen, 2017) or slow growth of the bacteria (Pamp *et al*., 2008).

But still, the classification of planktonic and biofilm growing bacteria is a complex task both experimentally and diagnostically. Evidently, we do not have other direct biomarkers than microscopic visualization to distinguish single cells and biofilms, which is impractical for routine purposes, such as in diagnostics. Here, we show that microcalorimetric measurements of metabolic energy release combined with a novel data analysis approach were able to differentiate the two growth forms.

Calorimetry has traditionally been used in materials research, using either isothermal titration calorimetry to determine, e.g. the Gibbs free energy (ΔG) of the binding of ligands to macromolecules or as differential scanning calorimetry to assess the stability of, e.g. proteins (Krell, 2008). In life sciences, isothermal microcalorimetry has been used to discriminate treatments or bacteria (Solokhina *et al*., 2017; Butini *et al*., 2019) and also to estimate minimum inhibitory concentrations (MIC) of antibiotics (Tellapragda *et al*., 2020). Simplistic measures such as maximum heat flow, total energy release, time to peak, etc. might be sufficient in some cases, but also ignore fundamental differences in the shape of the thermograms that may be related to characteristic metabolic processes.

### Metabolic differences between single cells and biofilms

The total energy in each well was not different between the models and is ultimately linked to either the availability of electron acceptors or -donors for respiration in the sealed well and we found total released energy similar to what was seen before (Braissant *et al*., 2015). Here we showed that the cease in metabolism was related to electron acceptor-rather than electron donor availability by supplementing with NO_3_^-^. *P. aeruginosa* is able to grow anaerobically with high biomass yield on NO_3_^-^ (Strohm *et al*., 2007; Line *et al*., 2014) and further on arginine and pyruvate fermentation (Eschbach *et al*., 2004; Schreiber *et al*., 2006) and reduction-oxidation reactions of self-produced phenazines (Price-Whelan *et al*., 2007). This supports the current setup for studying clinical biofilm where multiple studies have showed that bacterial metabolism is halted by electron acceptor availability (Kragh *et al*., 2014; Jensen *et al*., 2017) rather than carbon source.

We developed a numerical model that allows us to guess about the determining factors during growth. In this model, we chose to make it as simple as possible and only include two parameters (growth and O_2_ consumption) to see how much of the data could be explained by these simple factors. By making global fits where some parameters were kept constant between the growth-models (Table 1), the numerical model was able to explain a surprisingly high proportion of the thermograms by only fitting growth/die rates, initial concentration of bacteria, and a growth nonlinearity constant K_a_. All fits had R2 values of >0.9 and, not surprisingly, the fit for the planktonic culture was highest, since the model assumed a homogenous distribution of cells.

Such simple models cannot capture the complexity of the myriad of different metabolic pathways that *P. aeruginosa* can employ (Sønderholm *et al*., 2017a). Therefore, the secondary peak was not captured well by the model. This peak may be linked to a switch to alternative and lower yield metabolism as a response to the deprivation of O_2_ which is known to initiate the transcription of a suite of different genes (Guest, 1992) regulated by the concentration of different electron acceptors using e.g., FNR-type regulators (Unden and Schirawski, 1997). Additionally, the slow growth of biofilms is not necessarily equivalent to a low metabolic activity, as part of the metabolic energy will be used for maintenance and other growth unrelated processes (Kiviet *et al*., 2014; Monod, 1949; Sherman and Albus, 1924) not accounted for by this model. The numerical model suggested that *P. aeruginosa* grown in alginate beads was associated with higher death rates than the other models. For simplicity, we used the term death rate, but it might better be explained by a switch to an inactive state, as it has previously been shown that bacteria can resume growth after prolonged starvation and electron acceptor depletion (Kvich *et al*., 2019).

Interestingly, it seems that there are also fundamental metabolic differences between surface-attached and embedded biofilms. From this study, the mechanism by which they differ remains speculative but could be related to the differences in resource stratification experienced by bacteria in the different models. In the surface attached biofilm, a thin layer of cells was adherent to the bottom and sides of the well, and they will experience a less steep gradient of electron acceptor and carbon source. As a result of the more pronounced resource gradient in the alginate beads, a gradient of individual aggregate sizes is seen from the bead edge toward the center (Sønderholm *et al*., 2017b) with free space between aggregates. It has been shown that such zonation can create individual compartments with distinct pharmacokinetics (Christophersen *et al*., 2020) and we speculate that the microenvironmental characteristics could also cause distinct metabolic compartments (Kirketerp-Møller *et al*., 2020).

### Machine learning algorithm differentiates growth form

To analyze fundamental differences in the thermograms, we performed PCA analysis to investigate if clustering between groups existed (Fig. S2). From the PCA plot, it was evident that the three different growth forms had distinct signatures. This is a novel way of analyzing microcalorimetric data where the dimensionality of raw signals is reduced while minimizing information loss. To explore if we could use these signals as a potential “biofilm-biomarker” we designed an algorithm to analyze raw microcalorimeter signals. We used a warping function technique (Fig. S4) for correcting signals that vary in temporal dynamics, as taking the average of such signals will produce a smoothed-out version with none of the original features. In addition, we also used a warping function to correct for the difference in magnitude of heat flow signals. The warping in two dimensions allows for comparing more fundamental differences in the shape of the thermograms and not only use predefined parameters to describe differences or similarities with the risk of missing characteristic metabolic features.

By combining the dynamic warping approach and data augmentation, the variation of day to day experiments and small operator differences are widely imitated by producing variance by multiple iterations making the algorithm more robust. Resultantly, the algorithm was able to accurately classify each sample into the correct category for *P. aeruginosa*. The algorithm was designed using the thermograms of *P. aeruginosa* and was validated on *S. aureus* using the same classification categories into the specific growth-models. As these two microbes are distinct in both their metabolism, morphology and cell wall composition (gram positive vs negative), we did not expect them to fall into the exact same categories. However, there still seems to be a separation between the “biofilm-state” and single cell state for *S. aureus* except for the 24 h preincubated surface biofilm that clustered closer to the planktonic cultures. The algorithm was trained with a data point from the 24 h preincubated surface biofilm which resulted in the poor classification accuracy for the combined SB(24,48) category. This could have been accounted for by splitting this data into two separate categories. Similarly, the PC category overlaps with the SB category. From this study, it is unknown why the 24 h preincubated surface biofilm share more metabolic features with the planktonic culture but we speculate that *S. aureus* surface biofilms could need more time to change their metabolism into a “mature” biofilm phenotype than *P. aeruginosa*.

## Conclusions

Recently, it was shown that different bacterial species, organisms, and tissues displayed visually distinct thermograms (Braissant *et al*., 2015). Here, we propose a quantitative framework that could be used to classify different microbes and tissues based on their metabolic flux profiles. However, it is essential to note that the algorithm in its current form is a proof of concept that was developed using *in vitro* data. In e.g., infections, there will most likely be a mix of not only different species but also a mix of single cells and aggregates. The ability of our approach to differentiate such situations remains to be tested but we speculate that sophisticated analysis can successfully decipher such complex signals, similar to the process of spectral deconvolution.

Using our non-biased algorithm directly on the raw microcalorimetric data, we have elucidated that the specific heat flow of a bacterial culture reveals whether it is planktonic or biofilm growing which may be used as a biofilm biomarker. The biomarker can potentially be used as a diagnostic tool detecting the presence of different bacterial species and their growth state. Additionally, the analysis approach can also be used generally to classify differences in thermograms.

## Materials and methods

### Strains and growth conditions

We used a wild type *P. aeruginosa*, PAO1 strain obtained from the *Pseudomonas* Genetic Stock Center, ECU, USA (strain PAO0001) and a wild type *S. aureus* strain (NCTC8325) for microcalorimetric measurements. For microscopy, we used a PAO1 tagged with a stable green fluorescent protein (GFP) constitutively expressed by plasmid pMRP9 (Bjarnsholt *et al*., 2005). All experiments were performed at 37 °C in R2A broth (Lab M Ltd, UK) supplemented with 0.05 M Tris-HCl buffer (pH = 7.6) and 0.5 % glucose (henceforth mentioned as R2A media). Overnight (ON) cultures were started according to (Kragh *et al*., 2017) in R2A media.

### Preparation of planktonic cultures, surface-attached biofilms, and alginate beads

All samples were prepared inside plastic inserts (non-activated calWells, Symcel, Sweden). For surface-attached biofilms, the ON culture was diluted to a final optical density (OD_450_) of 0.005 with fresh R2A media. A 200 µL aliquot of diluted culture was inoculated into each insert, covered with parafilm, and incubated for either 24 or 48 hours at 37 °C, 120 rpm. This allowed biofilm to develop on the sides and bottom of the insert. Each insert was then washed with saline (0.9% NaCl) to remove planktonic biomass. After washing, 200 µL fresh R2A media was added and the insert was positioned in the calPlate (Symcel, Sweden).

For planktonic cultures, the ON culture was filtered through a sterile, syringe filter (ϕpore = 10µm) to remove aggregated bacteria. The filtered culture was then diluted to an OD_450_ of 0.005 with fresh R2A media. An aliquot of 200 µL was added to each insert and positioned in the calPlate (Symcel, Sweden).

Alginate beads containing bacteria were produced as previously described (Sønderholm *et al*., 2017b). Alginate beads were produced by mixing seaweed alginate (2% w/v) (Protanal LF 10/60 FT; FMC Biopolymer, Norway) with an ON culture adjusted to an OD_450_ of 2 to a final OD_450_ of 0.1. Beads were formed by extrusion dropping through a 21-gauge needle placed 3 cm above the surface of a stirred 0.25 M CaCl_2_ solution and left to harden for 1 h, producing beads of ϕ = 2.4 mm (Sønderholm *et al*., 2017b). Beads were rinsed in 0.9% saline and transferred to prewarmed (37 °C) R2A media. The beads were incubated in R2A media at 100 rpm at 37 °C for either 0, 24, or 48 hours. After incubation, beads were gently rinsed in 0.9 % saline to remove non-embedded cells from the bead surface. A single bead was then placed in each insert, the insert was filled with 190 µL fresh R2A media, resulting in a final volume of 200 µL in the insert. The insert was then positioned in the calPlate (Symcel, Sweden). Microcalorimetric measurements of *S. aureus* were performed identically to *P. aeruginosa*, but with Mueller-Hinton broth containing with 10mM KNO_3_ instead of R2A medium.

### Microcalorimeter procedure

Microcalorimetric measurements were conducted according to the manufacturer’s procedures and guidelines (Symcel, Sweden) and as previously described (Braissant *et al*., 2015; Wadsö *et al*., 2017). Each plastic insert was placed inside sterile, titanium cylinders with forceps. Each cylinder was sealed with a titanium lid and tightened to identical torque (40 cNm). A rack of 48 cylinders, including 32 samples and 16 references (filled with sterile media), was inserted into the microcalorimeter (calScreener, Symcel, Sweden) running the software calView 1.033. The rack was preheated in position 1 for 10 minutes, then moved to position 2 for 20 minutes before being moved into the measuring chamber. The wells were stationary during measurements and the system was allowed to equilibrate for approximately 30 minutes before stable signals were recorded. Measurements of heat flow (in µW) were recorded at a rate of 1 hertz.

### Numerical model

We constructed a system of coupled differential equations and solved them numerically to theoretical curves of metabolism, assuming that the metabolic behavior was non-linearly dependent on bacterial concentration and the O_2_ availability in the wells. For details, please see Supporting Information.

### Machine learning algorithm

We employed Gaussian Processes (Pedregosa *et al*., 2011) as the base machine learning algorithm. To augment single data points, we employed Gaussian warping functions. Briefly, we chose a small number M (M=5 here) and defined the warped signal of f(t), ∈ [0,T] as:

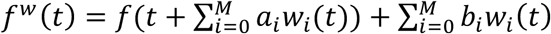

Where

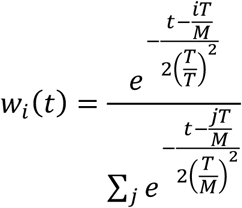

Here {a_i_} and {b_i_} are control points that determine smooth time shifts and smooth value shifts, respectively. These 2 × M values can be chosen to map one curve onto another as best as possible. In this sense, this is similar to a soft dynamic time warping with added value warping. See Supporting Information for details.

These warping functions were also used to generate augmented data for the training of the algorithm. In particular, we drew random numbers for {a_i_} and {b_i_}. The result of this is shown in Fig. S5.

## Statistics

ANOVAs and Tukey’s Multiple Comparison tests were performed in Prism (v. 7.0 GraphPad, USA). Principal component analysis was made in R (v. 3.6.3). Numerical model, and the machine learning algorithm was made in Python. CalScreener measurements were conducted with four technical replicates and four independent biological replicates from experiments conducted at four separate time points for *P. aeruginosa* and 4 technical replicates and three independent biological replicates from experiments conducted at three separate time points for *S. aureus*.

## Author contributions

ML, KK, and TB conceived and outlined the study, ML and KK performed experiments, ML, KK, BF, JK, and TB analyzed the data, JK performed mathematical simulations and made the machine learning algorithm, ML, KK, BF, JK and TB wrote the paper.

## Acknowledgments

This study was supported by the Lundbeck Foundation through grant R250-2017-633 (ML) and R105-A9791 (TB) and the Novo Nordic Foundation through grants NNF19OC0056411 and NNF19OC0054390 (TB). We thank Symcel for excellent technical assistance during the start-up of the project.

