## Supplemental information for "Metabolic flux fingerprinting differentiates planktonic and biofilm states of *Pseudomonas aeruginosa* and *Staphylococcus aureus*"

<u>Supporting Information</u>

Colony forming units (CFU) were quantified in parallel incubated samples at t0 and at the termination of the experiment. After the inserts were removed from the titanium cylinders biomass was collected by scraping with a pipette tip (Kragh *et al.*, 2019).

All liquid was then transferred from the wells to individual 1.5 mL Eppendorf tubes, as well as the alginate bead in the case of these samples. The alginate beads were dissolved in a 200 µl 0.02M/0.05M mix of citric acid and $Na_2CO_3$, respectively, followed by 10 min shaking at 1400 rpm which have been shown not to affect viability of the bacteria (Mater *et al.*, 1995). All samples were then degassed for 5 min followed by 5 min of ultra-sonication in an ultra-sound bath (230 VAC, Branson, USA) to break up aggregates. Subsequently, samples were 10-fold serial diluted and plated on LB plates (1.5% agar) and counted following incubation for 24 h at 37°C. CFU counts rely on two technical replicates in each three biological replicates.

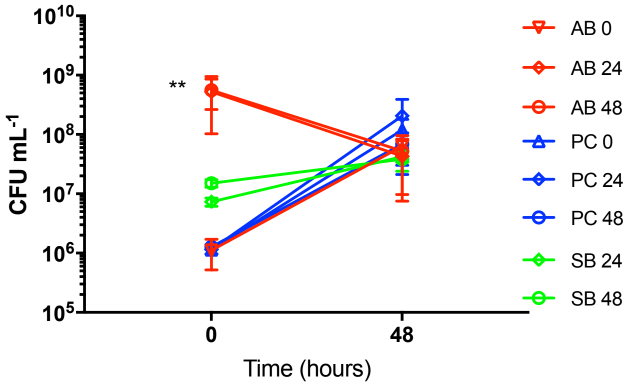


*Fig. S1. Development of number of bacteria in the different models from the samples were loaded into the calScreener (t=0) and ~48 hours later. The number of bacteria in the different models were statistically the same after 48 hours and at the start of the experiment ($p>0.05$; Tukeys multiple comparison test) except for the AB 24 and AB 48 at t=0 ($p<0.01$; Tukeys multiple comparison test).*

Numerical model of metabolic output

The calScreener measures the metabolic output of a collection of bacteria. We start by writing down a mathematical model of the behavior we expect in the simplest case of planktonic cells or a well-mixed suspension of cells.

Writing ρ(t) for the density of bacteria in a uniform suspension and c(t) for the oxygen concentration, we take

$$\partial_t \rho(t) = \alpha N_\rho(\rho) N_c(c) \rho - \beta_\rho$$

Here α is a growth rate and β a die rate. $N_\rho$ models the non-linear dependencies on the bacterial concentration, and $N_c$ the non-linear dependencies on the oxygen concentration.

The bacteria, growing initially exponentially, will fill out available space, creating a non-linearity in their growth. This we take to a general function of the form:

$$N_\rho(\rho) = \frac{1}{1 + (k_a\rho)^{\gamma\rho}}$$

Likewise, bacteria will stop growing once oxygen is gone. For this we take:

$$N_c(c) = \left(\frac{1}{2}[1 + \tanh(k_c c)]\right)^{\gamma c}$$

where $\gamma c$ accounts for any asymmetry in the function. Numerous choices could have been used for these functions, and most would work well. The key aspect is that some non-linearity is included.

The output of the calScreener is the metabolism M(t). The metabolism includes contributions from growth but also other functions involving energy. Thus, we have two terms, where only one of them depends on the non-linearity of growth:

$$M(t) = a[\in N_c(c)\rho + N_\rho(\rho)N_c\rho]$$

Finally, the oxygen is spent by the metabolism alone:

$$\partial_t c(t) = -\frac{1}{Y}[\in N_c(c)\rho + N_\rho(\rho)N_c(c)\rho]$$

We solve this coupled system of differential equations numerically to theoretical curves of metabolism (Fig. 3). We expect many of the parameters to be shared between the datasets, since we use the same species and strain of bacteria. The difference lies in the geometry of their growth and thus the only parameters we allow to vary between experiments is the effective growth parameters. In all cases we take c(t = 0) = 1, but allow b(t = 0) to vary. In other words: the initial oxygen concentration is the same, but initial bacterial concentration varies between experiments.

Principal component analysis

The raw heat flow values from all conditions and replicates from 0hr to 36hr where combined into a single matrix and analyzed by principal component analysis using the ‘prcomp’ function in R with default settings and no scaling. A PCA plot of each samples’ position along the first two principal component vectors was generated with the ‘factoextra’ and ‘ggplot2’ R packages.

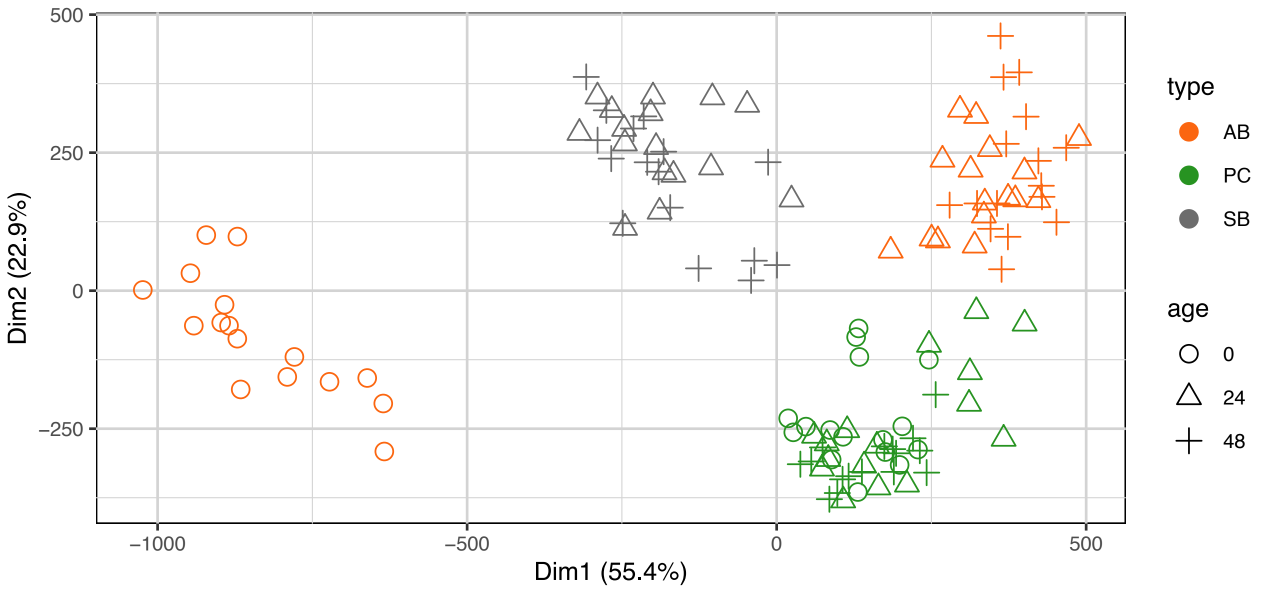


*Fig. S2. PCA of thermograms for P. aeruginosa grown in either alginate beads (AB), as planktonic culture (PC) or as surface biofilm (SB) preincubated for either 0h (circles), 24h (triangles) or 48 hours (plusses).*

Machine Learning

The time signals of repeat experiments vary a lot as shown in Fig. S4A. They vary in temporal dynamics: some metabolic curves peak faster than others, etc., and they vary in absolute numbers: some peak at higher values than others, etc. The variations could be caused by small day-to-day differences in e.g. sample handling etc. but are also affected by the number of bacteria loaded into each well of the microcalorimeter.

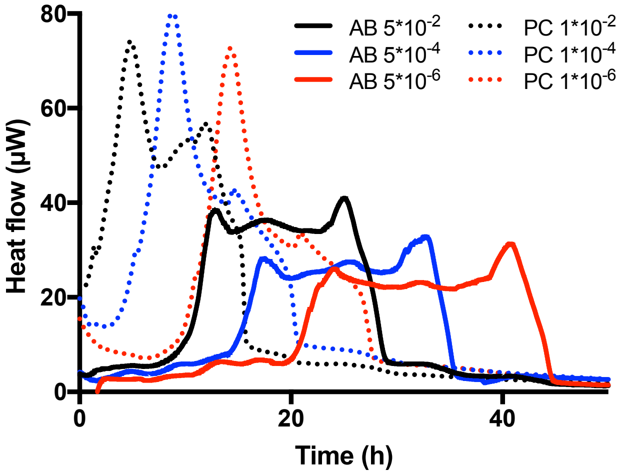

*Fig. S3. Effect on inoculum concentration on the position and magnitude of heat flow peaks. Solid lines represent bacteria inoculated into alginate beads (without preincubation) and dotted lines represent planktonic bacteria. Numbers in the legend refers to optical density ($OD_{450nm}$) values.*

To demonstrate this, we loaded bacteria in different dilutions into the wells (both as planktonic cultures and embedded in alginate beads) which caused both horizontal and lateral shifts in the position of peaks (Fig. S3). These variations complicate many operations. For instance, taking the average of the curves yields a smoothed-out version with none of the original features.
Dynamic time warping is a powerful technique for comparing signal that vary in speed. However, it does not handle variations in absolute numbers well. Instead here we introduce a simple warping function that allows us to map functions to lie closer towards one another. We choose a small number M (here we take M = 5) and defined the warped signal of f(t), t ∈ [0,T] as:

$$f^w(t) = f(t + \sum_{i=0}^{M} a_i w_i(t)) + \sum_{i=0}^{M} b_i w_i(t)$$

where

$$w_i(t) = \frac{e^{-(t-\frac{iT}{M})/2(\frac{T}{M})^2}}{\sum_j e^{-(t-\frac{jT}{M})/2(\frac{T}{M})^2}}$$

Here {ai} and {bi} are control points that determine smooth time shifts and smooth value shifts, respectively. These 2 × M values can be chosen to map one curve onto another as best as possible. In this sense this is similar to a soft dynamic time warping with added value warping. Fig. S4B shows the result applied to the AB(0) data.

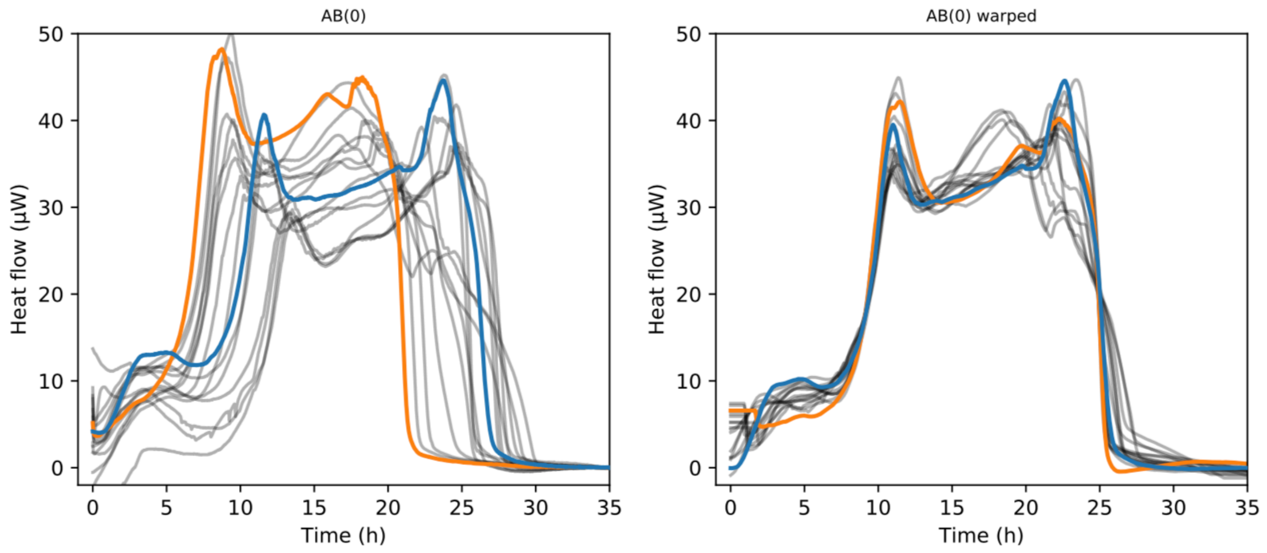


*Fig S4. A) Raw signals displaying variation in the position of peaks. B) warped signals in time and magnitude. Blue and orange lines are for eye guidance only.*

The warping function can also be used to generate augmented data. In particular, we can draw random numbers for {$a_i$} and {$b_i$} and do multiple iterations. The result of this augmentation is shown in Fig. S5.

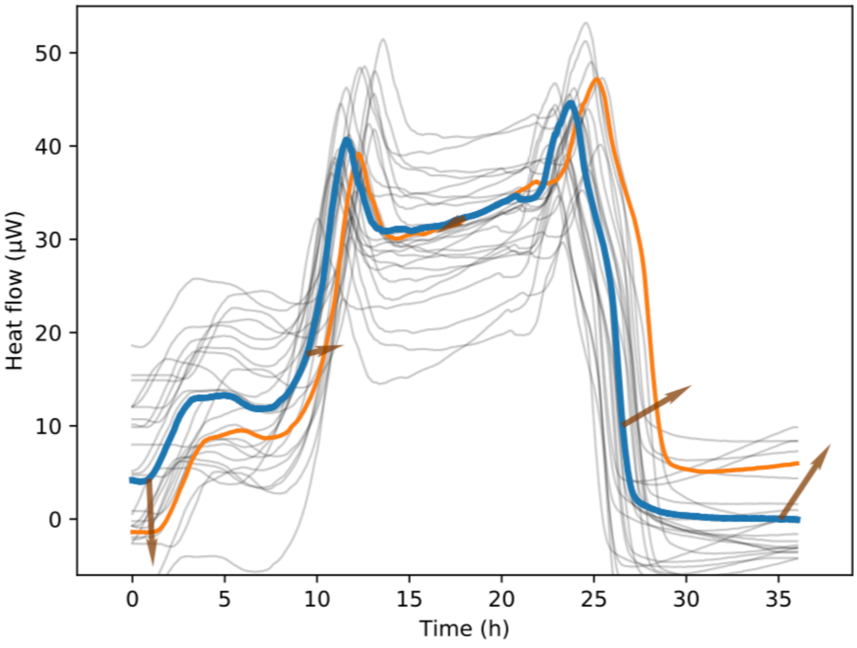


*Fig. S5. Augmented data created using warping functions. Blue and orange lines are for eye guidance only.*

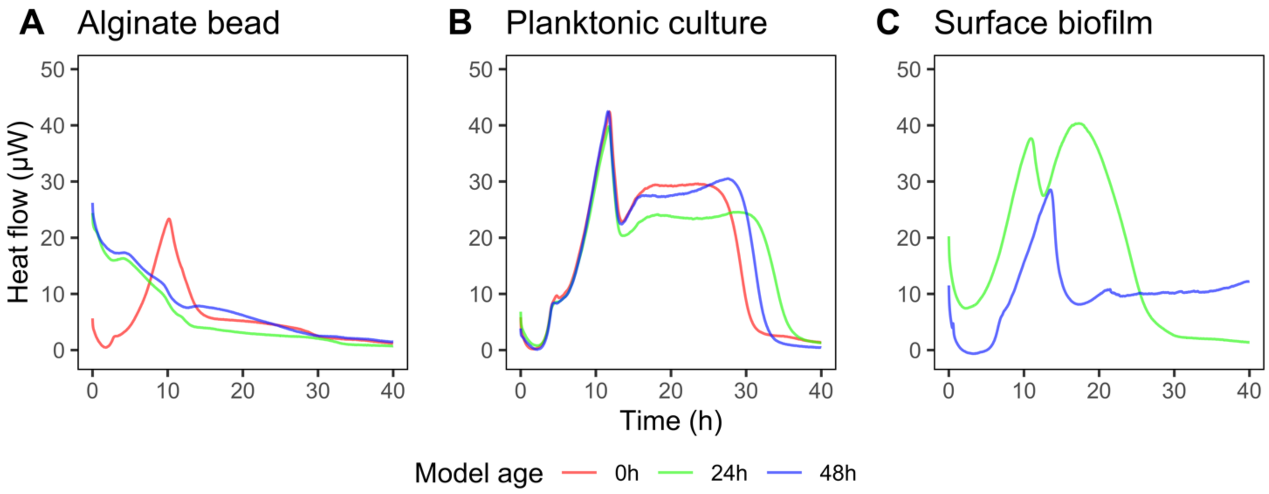


*Fig. S6. Example of thermograms for Staphylococcus aureus grown A) in alginate beads, B) as planktonic culture, and C) as a surface biofilm. Colors correspond to preincubation times of 0h (red), 24h (green) and 48h (blue).*
